# Epidermal stem cells self-renew upon neighboring differentiation

**DOI:** 10.1101/155408

**Authors:** Kailin R. Mesa, Kyogo Kawaguchi, David G. Gonzalez, Katie Cockburn, Jonathan Boucher, Tianchi Xin, Allon M. Klein, Valentina Greco

## Abstract

Many adult tissues are dynamically sustained by the rapid turnover of stem cells. Yet, how cell fates such as self-renewal and differentiation are orchestrated to achieve long-term homeostasis remains elusive. Studies utilizing clonal tracing experiments in multiple tissues have argued that while stem cell fate is balanced at the population level, individual cell fate - to divide or differentiate – is determined intrinsically by each cell seemingly at random ( 1 2 3 4 5). These studies leave open the question of how cell fates are regulated to achieve fate balance across the tissue. Stem cell fate choices could be made autonomously by each cell throughout the tissue or be the result of cell coordination ( 6 7). Here we developed a novel live tracking strategy that allowed recording of every division and differentiation event within a region of epidermis for a week. These measurements reveal that stem cell fates are not autonomous. Rather, direct neighbors undergo coupled opposite fate decisions. We further found a clear ordering of events, with self-renewal triggered by neighbor differentiation, but not vice-versa. Typically, around 1-2 days after cell delamination, a neighboring cell entered S/G2 phase and divided. Functional blocking of this local feedback showed that differentiation continues to occur in the absence of cell division, resulting in a rapid depletion of the epidermal stem cell pool. We thus demonstrate that the epidermis is maintained by nearest neighbor coordination of cell fates, rather than by asymmetric divisions or fine-tuned cell-autonomous stochastic fate choices. These findings establish differentiation-dependent division as a core feature of homeostatic control, and define the relevant time and length scales over which homeostasis is enforced in epithelial tissues.

---

For decades, research on tissue homeostasis has been focused on understanding the behaviors of resident stem cell populations for high turnover tissues ( 8 9). Recent work on stem cell populations in tissues such as the esophagus and skin epithelium suggests that stem cells are capable of stochastically undergoing both self-renewal and differentiation ( 1 2 3). Models that describe stem cell fate choices as cell-autonomous have been successful at describing the fate of individual cells *in vivo* with quantitative accuracy over time scales from days to years ( 2 3 4 5 10). Autonomous stochastic decisions have also been recapitulated in primary cultures ( 11). On the other hand, studies focusing on epithelial cell biology have described models where changes in cell density promote cell delamination ( 12), and division ( 13). Such models suggest a mechanism by which stem cells could coordinate their fates in homeostasis through ongoing density-dependent feedback, counter to the cell-autonomous view invoked in lineage tracing analyses. Although the existence of coordinated fate choice was shown in isolated crypts ( 14 15) and in specialized male germ line stem cells ( 16), its generality to other tissues, and how coupling occurs even in these systems, is not known. Thus, we sought to directly test the role of cell-cell coordination in promoting stem cell behavior and tissue homeostasis.

To study stem cell behaviors *in vivo*, we utilized the highly accessible mouse skin epidermis which provides a well-defined model to directly interrogate homeostatic stem cell behaviors over time ( 10 17 18). We and others have demonstrated that the epidermis contains a single stem cell population in its basal layer ( 2 3 10 19). Epidermal stem cell division is restricted to the basal layer, while differentiating cells delaminate from the basal layer and move into the suprabasal layer to contribute to the maintenance of the watertight skin barrier.

Existing strategies to study stem cell behavior rely on lineage tracing of spatially isolated cell clones. Although clonal fate data in the epidermis shows strong statistical signatures known as scaling behaviors, which are consistent with cell-autonomous fate models ( 2 3 4 5 6), it is less well appreciated that clonal statistics can appear to fit an autonomous model even when cells are strongly interacting. This result, recognized in statistical physics from the study of interacting systems known as Voter Models ( 6 7 20 21), is specifically predicted to confound single cell lineage tracing studies, but not the study of adjacent groups of cells. Therefore, to interrogate the existence of cell coordination in the epidermis, we required an alternative method that would track not only an isolated cell but all cells within a region of tissue.

We thus developed a novel lineage tracing approach, which does not rely on clonal labeling but rather on serial revisits of the same epidermal tissue through the utilization of the live imaging approaches developed in our lab (Fig. 1a). We further devised a semi-automated cell tracking method based on cortical and nuclear fluorescent epithelial reporters (*K14-GFPActin, K14-H2BCerulean*, respectively) to resolve individual cell behaviors across entire fields of view (Fig. 1b, Extended Data Fig. 1a,b). With this system, we tracked and identified the spatiotemporal distribution of basal cell self-renewal (cell division) and differentiation (basal layer delamination) events every 12 hours for seven days across a large region of epidermis. In total, we recorded 868 divisions and 849 differentiation events across three fields of cells from two mice. In addition, we recorded cell size, movement and density over time. This represents a comprehensive record of all events in each region.

**Figure 1.**
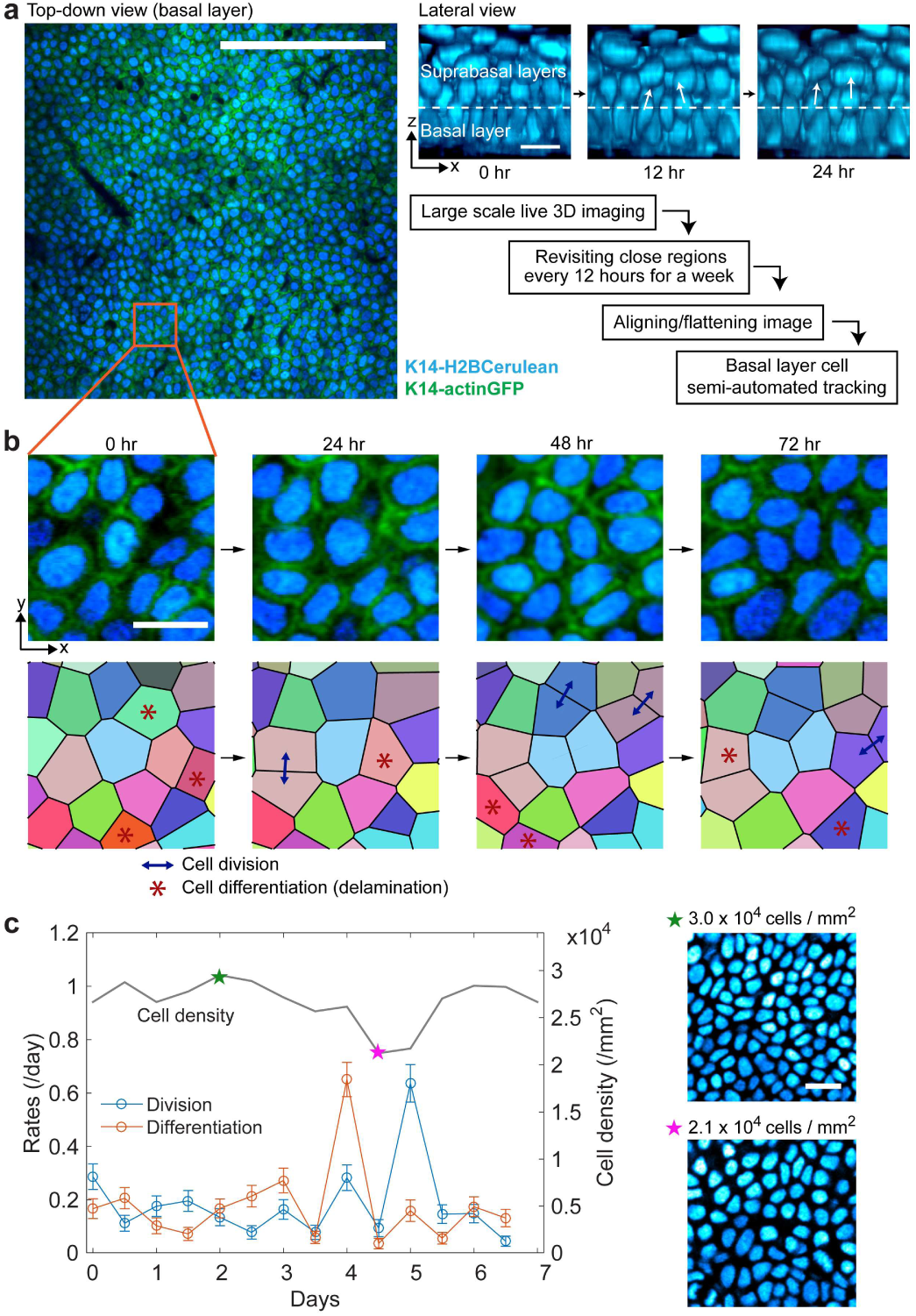
Serial revisits in mouse skin allows for live tracking of all epidermal stem cells in a region. **a,** Left: example of an optical section of the basal cell layer in a live mouse. Scale bar = 100 μm. Top right: lateral view of the epidermis with examples of cell differentiation (delamination from basal layer). Scale bar = 10 μm. Bottom right: schematic of experiment/analysis. **b,** Top: live tracing of an aligned region of the basal layer. Scale bar =10 μm. Bottom: voronoi diagrams delineating cells based on their nuclear signal (H2BCerulean). Colours represent clones from the initial timepoint; division/differentiation events denoted as shown. **c,** Left: time series of cell density, cell division rate, and cell differentiation rate obtained from a 90 μm x 90 μm region of basal layer. The density and rates fluctuate. Error bars represent sampling errors. Right: representative images of basal layer cells at high density (top) and low density (bottom). Scale bar = 10 μm.

Using this data, we looked for evidence distinguishing between two models of cell fate: cell-autonomous, as proposed previously in the epidermis ( 2 3 4 5 10), or through extrinsic regulation. In both cases, division and differentiation rates could be globally tuned by tissue-wide cues, and indeed we observed global fluctuations in these rates (Fig. 1c). To resolve between the two spatial models, we conducted a statistical test to quantify the time- and length-scales over which fate coordination occurs, if at all (Fig. 2a,b). Our test is based on the fact that cell-autonomous fate choices must give rise to stochastic imbalances in fate outcomes within a given region of tissue, akin to the imbalance in a coin flip between ‘heads’ (self-renewal) and ‘tails’ (differentiation). This imbalance, while averaging to zero in homeostasis, should nevertheless fluctuate in a mathematically predictable – binomial – manner over time and over extended regions of the tissue. In contrast, if cells coordinate their fates locally for instance, the number of self-renewal and differentiation events will be equalized much more precisely than binomial accuracy. We thus anticipated that if coordination occurs over a distance *l* and over a time *τ*, then the imbalance between the number of division and differentiation events would fluctuate as predicted by a cell-autonomous model for distances smaller than *l* and over time scales shorter than *t*, but these fluctuations would diminish in larger tissue patches over longer times (Fig. 2c).

**Figure 2.**
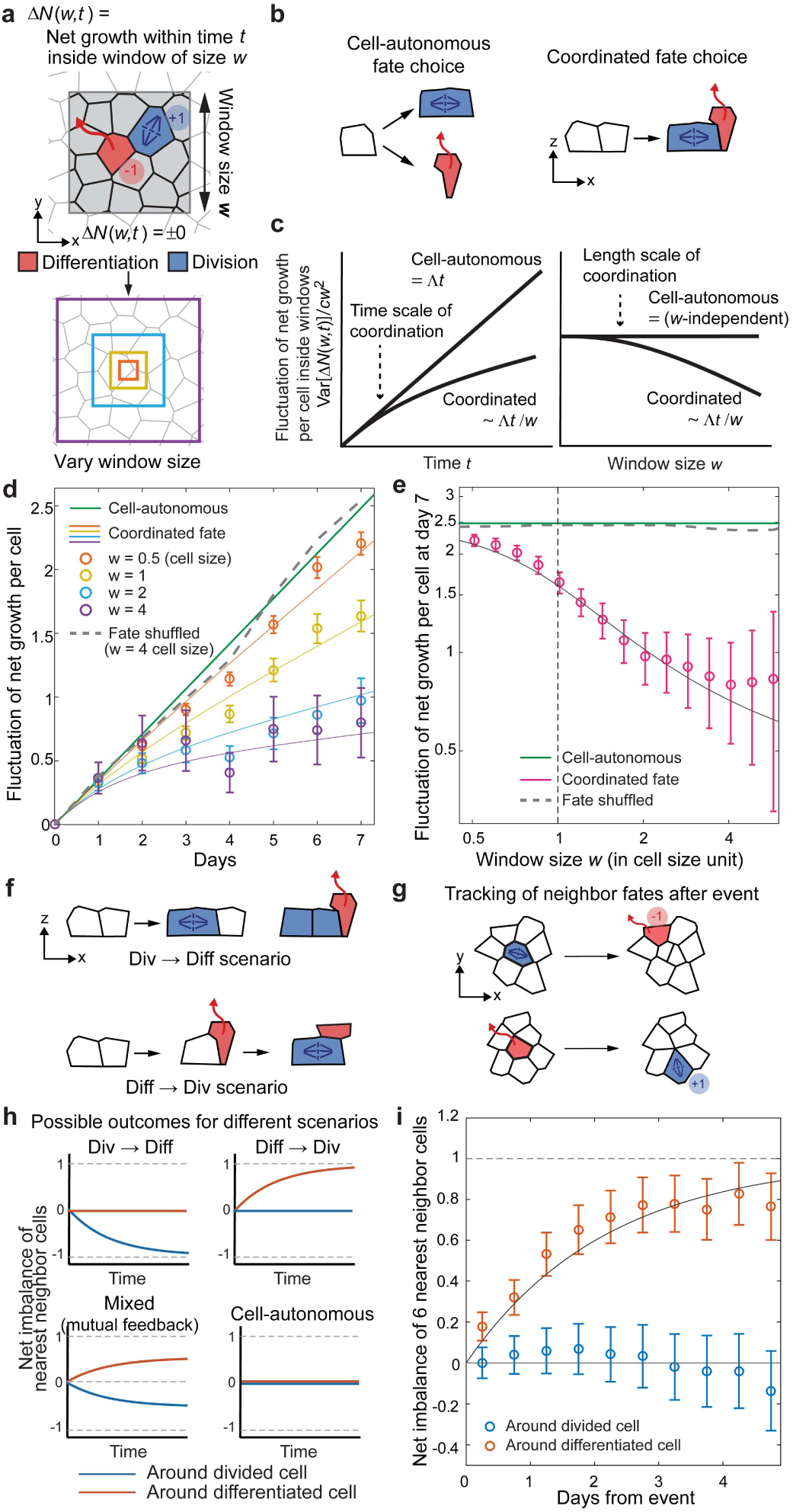
Differentiation and division are coordinated locally with temporal order. **a,** Schematic of the net growth fluctuation analysis. **b,** Cell-autonomous and coordinated fate choice models. **c,** Cell-autonomous and coordinated fate models predict different behaviors in the time evolution and window size dependence in the net growth fluctuation. c is the cell density and *Λ* is the fate choice event rate per cell. **d,** Time dependence of net growth fluctuation supports the coordinated fate model. Data from two separate 90 μm x 90 μm regions of same mouse, 523 divisions and 528 differentiations. Solid lines: model with the coordination time scale τ = 2.1 days and length scale *l* = 3.9 μm. Error bars: s.e.m. from bootstrapping. See Methods for the detail of theory and analyses. **e,** Window size dependence of net growth fluctuation. Same data, model, and error bars from **d. f,** Examples of possible ordering of events in the coordinated fates model. g, Schematic of the neighbor net imbalance analysis to resolve the time ordering. In each fate choice event, we subsequently tracked the six neighboring cells and quantified their net imbalance between division and differentiation. **h**, Predicted outcomes of the neighbor net imbalance analysis for different scenarios. **i,** Quantification of the net imbalance of six nearest neighbor cells (based on distance) shows that the differentiation precedes division, within a short time scale. Solid line: 1-exp(-*t*/*τ*), with *τ*=2.1 days. Error bars: s.e.m. of primary divisions and differentiation events.

To formalize this insight, we defined the imbalance between the number of division and differentiation events as a function of window size w and time *t, ΔN*(*w,t*), and predicted the fluctuations for the two alternative hypotheses (see Methods for detail of the theory). Comparing the predicted fluctuations to our data, we found strong evidence for the presence of cell-cell coordination in fate choices (Fig. 2d,e, Extended Data Fig. 2a,b,c). By fitting to a model of spatio-temporal fate coordination, we found that the length scale for cell fate coordination was comparable to the cell radius (~4 μm), suggestive of direct nearest-neighbor interactions.

We next sought to interrogate the sequence of self-renewal and differentiation events between neighboring cells over time (Fig. 2f). If cell division induces differentiation of neighboring cells, we would expect to see a net imbalance toward differentiation adjacent to dividing cells, whereas if differentiation induces nearby cell division, a reciprocal increase in division would be expected (Fig. 2g,h). Additionally, both forms of feedback could be present, whereas if division and differentiation were uncoupled, we would expect no neighboring behavior compensation. We found a clear unidirectional bias, with one net additional stem cell division following a neighboring cell differentiation event, but no reciprocal compensation (Fig. 2i). This behavioral sequence bias is fast acting (~2 days) and occurs only at short range (nearest neighbor, Extended Data Fig. 3a), consistent with our fluctuation test (Fig. 2d,e). Thus, these results provide a simple and robust model for locally enforcing tissue homeostasis: stem cell differentiation (basal layer delamination) is directly followed by division of a neighboring stem cell to balance epidermal cell density.

**Figure 3.**
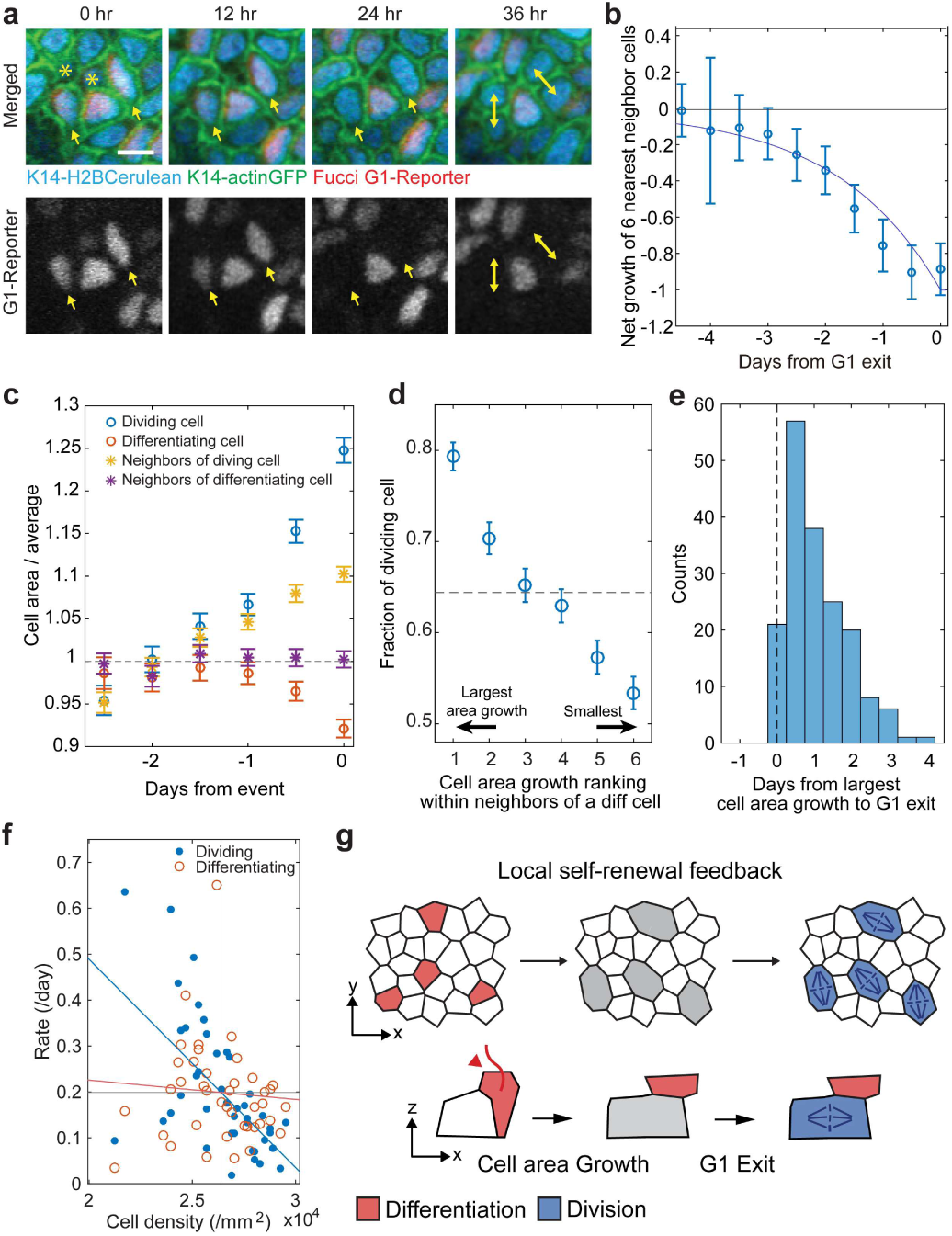
Differentiation triggers local density drop which induces division commitment in neighboring cell. **a,** Illustrative time series of a Fucci G1-reporter in epidermal stem cells. Arrows show two cells losing fluorescence after exiting G1 and progressing to division (double headed arrows) following neighbor cell delamination (indicated by *). Scale bar = 10 μm. **b,** Accumulated net imbalance of division and differentiation events surrounding a dividing cell at times preceding G1 exit, showing cell delamination reliably occurring between 1-3 days prior to S phase entry (N=187 division events from two separate imaging regions). **c,** Dynamics of the average area of cells in the time leading up to division/differentiation, as well as the average area of the cells in the nearest neighborhood. Error bars represent s.e.m. (N=868 divisions and 849 differentiations from 2 mice. **d,** The probability of a cell to commit to divide depends strongly on its fold-change in area following neighbor cell differentiation. Error bars represent sampling error (N=849 differentiation events). **e,** Histogram of the time delay between peak area fold-change and G1 exit; The delay is always positive or too short to observe, showing that cell expansion always precedes G1 exit (N=187 divisions). **f,** Correlation between average cell density across the entire region of observation (90 μm x 90 μm), and division/differentiation rates. Division rate was negatively correlated with density (R=-0.60, p<0.001), but the differentiation rate was not (n.s.). N=3 regions from two mice. **g,** Schematic of the proposed ordering of events in skin epidermal homeostasis.

To better understand how division locally follows differentiation, we decided to investigate cell cycle progression of stem cells subsequent to neighboring differentiation events. We generated mouse lines that contained epithelial fluorescent markers in addition to a Fucci G1-reporter (hCdt1-mKO1) ( 22) which specifically accumulates fluorescent signal during G1 of the cell cycle and drops signal in progression from G1 to S/G2 (Fig. 3a, Extended Data Fig. 3b). By retrospectively tracking dividing cells we found bias in neighboring differentiation events preceding S phase commitment, (Fig. 3b), supporting a model where differentiation facilitates G1-to-S phase cell-cycle progression and subsequent division of neighboring stem cells.

Previous studies have suggested that changes in epithelial cell density can drive cell division ( 12) and delamination ( 13). Therefore, we next asked whether the observed coordination of cell fates could be mediated indirectly by the degree of crowding experienced locally by cells. By estimating the average effective cell area prior to cell division or differentiation, we observed that dividing cells began to expand their basal footprint ~1.5 days prior to division, while differentiating cell basal area shrank (Fig. 3c). Notably, the cell density in the neighborhood of a dividing cell dropped prior to division, but no density change was observed in neighboring cells preceding a differentiation event (Fig. 3c). Moreover, following a differentiation event, the neighboring cell showing the greatest fractional increase in area had the highest probability to divide (Fig. 3d). We further found that the G1 phase exit in a dividing cell always happened at the same frame or after basal area increase (Fig. 3e). The local density responses were also reflected in the global tissue dynamics: we found that the global cell density negatively correlated with the cell division rate, but not with the cell differentiation rate (Fig. 3f). Thus, a drop in basal cell density precedes cell division commitment (Fig. 3g), but overcrowding does not precede delamination during homeostasis.

Our analyses suggest that epidermal stem cell delamination drives the division of neighbors, but not the opposite. To functionally test this ordering of events, we asked how inhibiting cell division affects basal layer dynamics. We predicted that if cell delamination is unaffected by cell density or proliferation it should continue even when stem cell divisions are blocked, leading to a rapid drop in density in the basal layer (Fig. 4a). By inhibiting cell division using Mitomycin C (MMC) or Democolcine ( 23), we found indeed that the cell density progressively reduces in the basal layer, but not in the suprabasal layers when compared to vehicle control (Fig. 4b,c; Extended Data Fig. 3a,b). Cell tracking combined with inhibition of cell divisions showed that differentiation from the basal layer is largely unaffected when epidermal cell divisions are blocked and can persist for multiple days without epidermal cell divisions (Fig. 4d,e).

**Figure 4.**
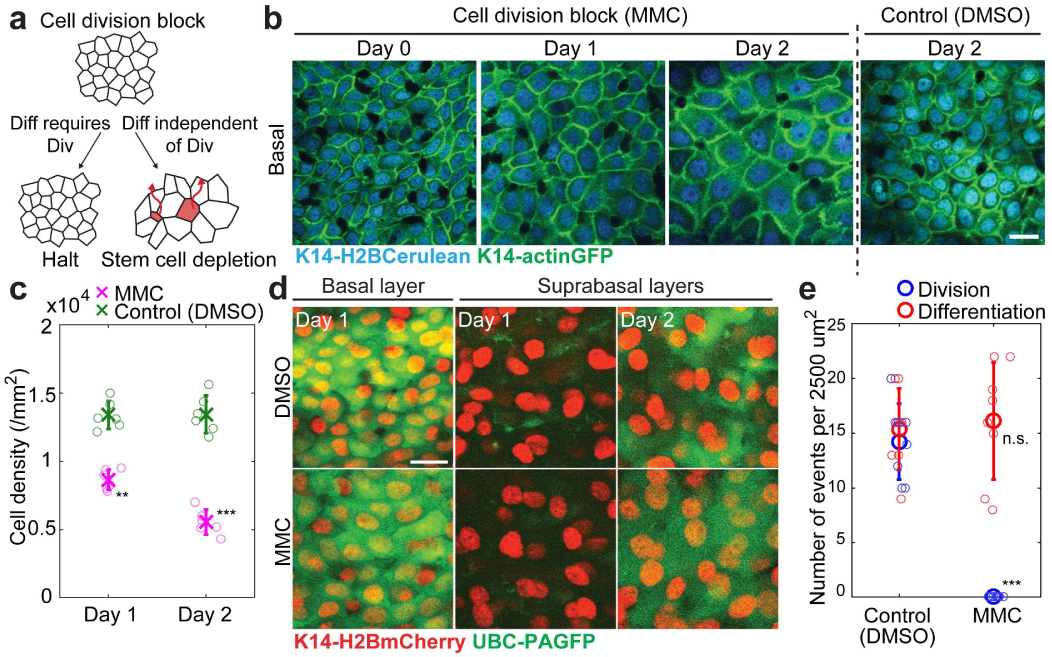
Differentiation is independent from division. **a,** Schematic of possible outcomes in the basal layer following inhibition of epidermal proliferation. Red indicates differentiating cell. **b,** Inhibition of stem cell divisions through Mitomyocin-C (MMC) treatment leads to progressive reduction in basal cell density. Scale bar = 10 μm. **c,** Epidermal basal density. Error bars represent s.d. (N=6 regions from 2 mice). **d,** Photolabeled cells traced during inhibition of cell division. Cell density in the basal layer decreased while no apparent change occurred in the suparabasal layers. **e,** Basal cell behaviors with and without MMC. Error bars represent s.d. (N=8 regions from 2 mice).

Altogether, we have found that sustained epidermal tissue homeostasis relies on a local feedback mechanism whereby differentiating stem cells that delaminate from the basal layer lead to the G1 exit and subsequent self-renewal of neighboring stem cells. At the level of individual cells, and even clones, cell fate was previously observed to be stochastic; only by studying the spatial and temporal ordering of cell behaviors does it become apparent that fate choices are in fact tightly and directionally coordinated, providing a robust mechanism for balancing stem cell behaviors while allowing a rapid and sustained response to fluctuating tissue demand. Understanding how this homeostatic program breaks down in hyper-proliferative disease states, such as cancer, will be fundamental for future therapeutic interventions.

## Acknowledgments

We thank Elaine Fuchs for *K14-actinGFP* mice. This work is supported by The New York Stem Cell Foundation and grants to V.G. by the Edward Mallinckrodt Jr., an HHMI Scholar award and the National Institute of Arthritis and Musculoskeletal and Skin Disease (NIAMS), NIH, grant no. 5R01AR063663-04; grant no. 1R01AR067755-01A1. NIH Predoctoral Program in Cellular and Molecular Biology (K.R.M. grant T32GM007223). The content is the responsibility of the authors and does not necessarily represent the official views of the NIH. K.R.M was an NSF Graduate Research Fellow. V.G. is a New York Stem Cell Foundation Robertson Investigator and HHMI Scholar. A.M.K. is supported by a Career Award at the Scientific Interface from the Burroughs Wellcome Fund, and an Edward Mallinckrodt Jr. Foundation Grant. K.K. acknowledges Grants-in-Aid for JSPS Fellows (28-908). K.C. is supported by a Canadian Institutes of Health Research Postdoctoral Fellowship.

## Author contributions

K.R.M., K.K., A.M.K. and V.G. designed the project. K.R.M. and V.G. designed experiments and wrote the manuscript. K.K. and A.M.K. performed data analysis, statistical modeling and wrote the manuscript. K.R.M., D.G. and K.C. performed the experiments and analyzed the data. T.X. generated the K14-H2BCerulean mouse. J.B. assisted with technical aspects.

## METHODS

### Transgenic mice

K14-H2BCherry and K14-H2BCerulean mice were generated by the Yale Transgenic Facility. K14-actinGFP mice were obtained from E. Fuchs. Bsi596mKO2 (Fucci G1-reporter) mice were developed by Dr. Asako Sakaue-Sawano and Dr. Atsushi Miyawaki (RIKEN). UBC-PA-GFP mice were obtained from Jackson Laboratories. All procedures involving animal subjects were performed under the approval of the Institutional Animal Care and Use Committee (IACUC) of the Yale School of Medicine.

### Experimental treatment of mice

Preparation of both mouse plantar and ear skin for intravital imaging was performed as described recently ( 10 17 18). Briefly, mice were anesthetized with IP injection of ketamine/xylazine (15 mg/ml and 1 mg/ml, respectively in PBS). The ear epidermal areas were shaved using an electrical shaver and depilatory cream (Nair). After marking the area to be imaged for subsequent identification with a microtattoo, mice where returned to their housing facility. For subsequent revisits the same mice were processed again with injectable anesthesia. The plantar and ear epidermal regions were briefly cleaned with PBS pH 7.2, mounted on a custom-made stage and a glass coverslip was placed directly against the skin. Anesthesia was maintained throughout the course of the experiment with vaporized isofluorane delivered by a nose cone.

### Topical drug treatment

To inhibit cell proliferation in the epidermis, Mitomycin C (MMC) and Democolcine(23) were both delivered topically by applying it to ear skin. MMC was dissolved in a 15 mg/ml stock solution in dimethyl sulfoxide (DMSO), while Democolcine was dissolved in a 25 mg/ml stock solution in the DMSO. The stock solution was diluted 100 times in 100% petroleum jelly (Vaseline; final concentration is 150 μg/ml). One hundred micrograms of either working concentration were spread evenly on the skin area daily. A mixture of 100% DMSO in petroleum jelly (1:100) was used as a vehicle control.

In the analyses of the treated regions, the stars in Fig. 4c,e and Extended Data Fig. 4b represent the statistical significance (**,***,****, and n.s. representing *p* < 0.01,0.001.0.0001, and nonsignificant, respectively).

### *In vivo* imaging

Image stacks were acquired with a LaVision TriM Scope II (LaVision Biotec, Germany) microscope equipped with both Chameleon Vision II and Discovery (Coherent, USA) 2-Photon lasers. For collection of serial optical sections, a laser beam (940nm for GFP/Cerulean and 1100nm for mCherry, respectively) was focused through a 20X or 40X water immersion lens (Zeiss W-Plan-APOCHROMAT, N.A. 1.0; Zeiss W-LD C-APOCHROMAT, N.A. 1.1 Zeiss) and scanned with a field of view of 0.5 mm x 0.5 mm at 600Hz. z-stacks were acquired in 1 μm steps for a ~40-80 μm range, covering the entire thickness of the epidermis. Cell tracking analysis was performed by re-visiting the same area of the epidermis in separate imaging experiments, as described in Image Analysis. A micro-tattoo was introduced in addition to using inherent landmarks of the skin to navigate back to the original region; including the vasculature and distinctive clustering of hair follicles.

### Photo-activation

Photo-activation in UBC-PA-GFP mice was carried out with the same optics as used for acquisition. An 810 nm laser beam was used to scan the target area. Activation of the PA-GFP was achieved using 3% laser power for 30 sec.

### Image analysis

Images which included the suprabasal layer, basal layer, and the extracellular matrix were obtained as large tiled image stacks at roughly the same positions every 12 hours for 7 days guided by the microtattoo. We first manually aligned the images over the time course in Imaris (Bitplane) by using data from all three channels: K14-actinGFP, K14-H2BCerulean, and Fucci G1.

For the cell-tracking analysis performed by MATLAB scripts, we first cropped out three regions with size 115 um x 115 um which typically included more than 300 basal layer cells each. To correct for the difference of height positions of the basal layer within the 3D images, we first Gaussian blurred the signal from the K14-actinGFP channel spatially in the *xy*-plane (width 4 um) to create a 3D mask representing the region covering the whole epidermis (Extended Data Fig. 1a). We then defined the height of the interface between the epidermis and the dermis from the 3D mask, and subtracted this height from the original 3D data to level the basal layer position (Extended Data Fig. 1b).

From the height-corrected 3D images, we took a single z-position containing the nucleus of all the basal layer cells (Extended Data Fig. 1b), and performed cell tracking in 2D over the time-course. For this, we first manually corrected the shifts in the 2D images to minimize the overall depositions of cells, and then ran an automatic cell tracking algorithm based on the 2D positions of local maxima in the H2BCerulean channel. The algorithm assigned each cell (represented by the local maxima, calculated after Gaussian blurring the 2D image with width 1 μm) to a closest cell in the previous time frame. Tracked cells were frequently lost or were associated with more than one cell in the subsequent time frame, which indicated cell differentiation (i.e., delamination from the basal layer) and cell division, respectively. After manually correcting the errors in the tracking with guide from the height-corrected 3D images in all three channels, the script outputted the positions of the local maxima in the H2BCerulean channel and the lineages of the cells present in the basal layer at each time point.

### Quantification of cell area and G1 reporter signal level

At each time frame, we created a voronoi diagram based on the cell positions (=local maxima of H2BCerulean channel). The areas of voronoi regions associated to cells were recorded as the cell areas.

For the Fucci G1-reporter channel, we first z-score normalized the signal using all pixels at each time frame to account for the temporal fluctuation. Using this corrected signal, we calculated the signal per area of the cells at each time point using the voronoi diagram. We subtracted the background signal, defined as the average of the 10 cells with smallest signal per area at each time point.

Extended Data Fig. 3b shows the traces of the signal/area values for dividing cells (after subtracting background). Not all the cells had strong Fucci signal, and many of them had a low signal throughout their lifetime. Nevertheless, by selecting the cells based on the maximum level of signal/area (threshold set to include 187 out of 581 dividing cells across two regions), we found the expected time courses of the G1-reporter: signal going up and rapidly dropping before cell division. For later convenience, we define *s_f_*(*i*)=1 or 0 as the function telling if the cell labeled as *i* is a selected cell with respect to this criterion of G1-reporter (=1) or not (=0).

For the selected cells *s_f_*(*i*)=1, we defined the timing of G1 exit *t_d_*(*i*) as the time where the cell experienced the largest fold change decrease in the signal/area value. We observed that all the drops in G1 signal happened right before or 1 frame before the cell division, indicating that after G1 exit the cell divides within 24 hours.

### Fluctuation analysis

In the lineage tracing of the basal layer, the *xyt*-coordinate of a cell differentiation was identified as the cell position and the time frame right before the cell delaminated. Similarly, the *xyt*-coordinate of a cell division was determined as the cell position and time frame right before the cell underwent cytokinesis into two daughters.

Using these *xyt*-coordinates of the differentiation and division events, we conducted a fluctuation analysis by (1) randomly choosing 4096 positions of a window with size *w* within each region, (2) calculating the time course *ΔN*(*w, t*) for each sampled window as defined in the text, and (3) calculated the variance of *ΔN*(*w, t*) for each *w* and *t*. To avoid the tracking errors occurring near the boundary of the images, the windows were sampled from a fixed central region with size 90 μm x 90 μm. To generate the shuffled control (Fig. 2 d,e), we used the same *xyt*-coordinates but with randomly permuted fates (differentiation or division). The final ‘shuffled’ curves in Fig. 2d,e are the average of 128 independent randomizations of the data set.

The reported variances were then corrected for a bias resulting from window overlap. Randomly positioning windows with size *w* in a finite image region of size *L* will lead to overlaps. The overlap affects the number of independent windows that contribute to *ΔN*(*w, t*), and will effectively decrease the fluctuation. When randomly putting two windows inside a region of (2*w* ≤ *L*), the expected proportion of overlap in the area is

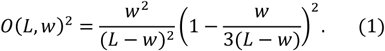

We divided the variances calculated as described above by 1 – *0*(*L,w*) to generate the corrected variance Var [Δ*N*(*w*, *t*)] plotted in Figs. 2d,e.

Equation (1) is derived as follows. Randomly placing a line segment with length *w* to fit inside the region with length *L*(≥ *w*) is equivalent to randomly sampling the left edge of the segment from [0, *L* – *w*]. Taking two random positions *x*_1_ and *x*_2_ from [0, *L* – *w*], the probability distribution of the distance between *x*_1_ and *x*_2_, *Δ_x_* = |*x*_1_ – *x*_2_|, is *P*(*Δ_x_*) = (*L – w – Δ_x_*)/(*L – w*)^2^ for 0 ≤ *Δ_x_* ≤ *L – w* and zero otherwise. The overlapping fraction of the two segments with the left edges positioned at *x*_1_ and *x*_2_ is 1 – *Δ_x_/*w for 0 ≤ *Δ_x_* ≤ *w* and zero otherwise. We next introduce a second direction *y* to randomly place line segments with bottom edges placed at *y*_1_ and *y*_2_. We then have two windows with the bottom left corners with the coordinates (*x_1_,y_1_*) and (*x_2_*,*y*_2_). Assuming that *L* – *w* ≥ *w* (*2w* ≤ *L*), the average overlapping fraction of the two windows is then

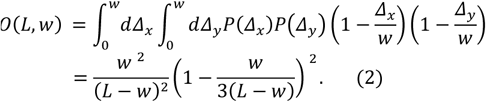

### Theory

Here we describe the theoretical model used in the fittings presented in Fig. 2d,e. We assume that the fate choice events, differentiation or division, will all happen in a coordinated manner but with stochastic time difference *s* > 0 and *xy*-displacement (*ξ*, *η*). We assume for simplicity that *s* is sampled from an exponential distribution: *P*(*s*) = *e*^−*s/r*^/*τ*, and (*ξ*, *η*) is sampled from a Gaussian distribution: 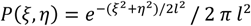. Here, *τ* and *l* are the time scale and length scale of the coordination, respectively, which are the fitting parameters. We also assume that *s* and (*ξ*, *η*) are independent from each other. This simple model is used to fit the key features of the data and to gain an intuition for the effect of lags and coordination distance on the observed behaviors, and is not intended to reflect any specific mechanism of fate coupling.

We consider a three-dimensional box with width and height of the window size *w* and the depth of the time length *t*. Sampling the pairs of events randomly across space and time, and assigning the above probability for the time difference and the displacement between the fate-coordinated pairs, we count the divisions that happened inside this box as +1 and the differentiations as -1, corresponding to cell increase and decrease events, respectively. We are interested in the statistics of the net growth, *ΔN*(*w*, *t*), which is the sum of the +1 and -1’s that occurred inside the box.

Since the contributions from the fate-coordinated events are zero (one differentiation and one division amounts to ±0), and since the isolated events (events inside the box that have a fate-coordinated pair outside the box) are binomially distributed between division (+1) and differentiation (-1) fates, the variance of *ΔN*(*w*, *t*) is equal to the average number of isolated events. Given that one of the events is sitting inside the box, the probability of the fate-coordinated event to be also sitting inside the box is

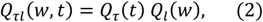

where *Q_τ_* (*t*) is the probability of finding the fate-coordinated event inside the time frame *t*, and *Q_l_*(*w*) is the probability of finding the fate-coordinated event inside the window of size *w.*

The average number of total events inside the box is *Λw*^2^*t*, where *Λ* = *c* (*λ* + *Γ*) is the rate per area of events with *λ*, *Γ*, and *c* being the steady-state cell division rate, differentiation rate, and cell density. The average number of isolated events, and thus the variance of *ΔN*(*w*, *t*), is obtained as

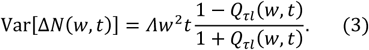

Here, the factor arises from the ratio of the isolated vs coordinated cells in the total of events, 1 – *Q_τl_*(*w*, *t*): 2*Q_τi_*(*w*, *t*). The factor 2 here arises from the pairs of cells that are both inside the box contributing as 2 cells in the total number.

We first compute *Q_τ_*(*t*). Assuming that the time difference between the two events is *s*, the probability of finding both events within the time frame *t* under the condition that at least one is inside, is 1 − *s/t*. Taking the average of this probability over *s*, we have

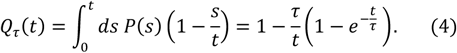

Note that the integral is only taken up to t since for pairs of events which are further apart from *t*, there is no possibility of finding both events inside the interval *t.*

Similarly, by assuming that the *xy* displacement between the two events is (*ξ*, *η*), the probability of finding both events fitting inside the window of size *w* under the condition that at least one is inside, is (1 – |*ξ*|/*w*)(1 – |*η*|/*w*). Taking the average of this probability over (*ξ*, *η*), we have

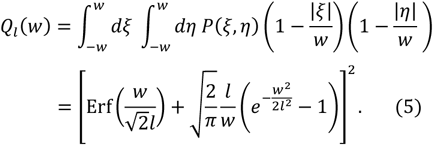

Here, 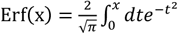 is the error function.

To understand how the coordination in fate affects Var[Δ*N*(*w*, *t*)], we consider two limiting cases: *l*/*w*, *τ*/*t* → *∞* and *l*/*w*, *τ/t →* 0. In the first case, since the fate-coordinated pairs are so far apart in time and space compared with *t* and *w*, the fluctuation should be equivalent to the case of the cell-autonomous model. Indeed, *Q_τ_*(*t*),*Q_l_*(*w*) → 0 in this limit, meaning that

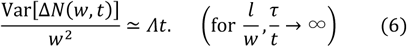

which is the statistics of the cell-autonomous model. In the second case, since *t* and *w* are much bigger than the coordination time and length scales, the isolated events can only be found near the surface of the box, meaning that the fluctuation should be one order smaller than the cell autonomous case in terms of *t* or *w*. We obtain *Q_τ_*(*t*) ≃ 1− *τ*/*t*, and 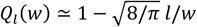 in the lowest order of *l*/*w* and *τ*/*t*, which leads to

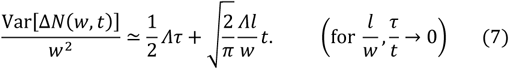

Here, the first term is a constant that does not depend on *t*, which is the contribution from the isolated events close to the initial timepoint or the final timepoint. The second term is linear in time but inversely proportional to *w*, which is the contribution from the isolated events sitting close to the edge of the window.

In Fig. 2 d,e, Extended Data Fig. 2a,b, we show the best fit of Eq. (3) to the data of Var[*ΔN*(*w*, *t*)] obtained by the least-squares method while using *Λ* = 9.4×10^3^ events day^-1^ mm^-2^ for the first mouse (Fig. 2d,e) and *Λ* = 1.3×10^4^ events day^-1^ mm^-2^ for the second mouse (Extended Data Fig. 2a,b). The best fit parameters were *τ* = 2.1 days and *l* = 3.9 μm (Fig. 2d,e) and *τ* = 1.0 day and *l* = 4.9 μm (Extended Data Fig. 2a,b).

### Forward tracking of neighbor fate imbalance

For each cell, labeled by *i*, there is the birth time *t_b_*(*i*), the fate choice time *t_f_*(*i*), and the fate *σ*(*i*) = ±1. If the cell was present at the initial time point *t* = 0, we set *t_b_*(*i*) = − *∞*. If the fate was not chosen before *t*_max_, which is the last time point of the image sequence, we set *t_f_*(*i*)= *∞* and *σ*(*i*) = NaN. Let us define the time course of imbalance for *t_b_*(*i*) ≤ *t* < *t_mbx_*:

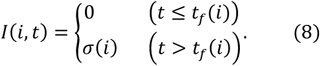

For *t* outside of this defined region, *I*(*i*, *t*) returns NaN.

We denote the label of the *k*-th nearest neighbor of cell *i* (in terms of *xy*-coordinate) at time t as *NN*(*i*, *t*, *k*). For each fate decision event of cell *i*, we calculated the net imbalance of the *K*-nearest neighbor cells in the subsequent time course:

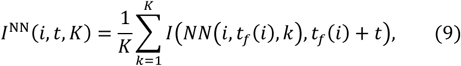

where *t_f_*(*i*) ≤ *t* < *t*_max_. Again, for *t* outside of this defined region, *I*^NN^(*i*, *t*, *K*) returns NaN. The sum in the right-hand side skips entries with NaN values.

To obtain the background imbalance, we first define *N*(*t*) as the number of cells that existed at time *t*, and *C* (*t*,*j*) as the label of the *j*-th cell at time *t* (1 ≤ *j* ≤ *N*(*t*)). The background imbalance within the imaged region is calculated as

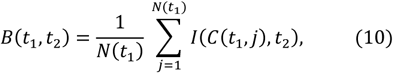

for *t*_1_ < *t*_2_. Note that *B*(*t*_1_,*t*_2_) ≠ 0 even in the ideal case (i.e., infinite number of samples and time constant rates). This is because the number of dividing cells vs differentiating cells within a randomly selected population is not 0.5 when the average lifetimes of dividing cells and differentiating cells are different.

By subtracting the background imbalance, we calculated the de-trended net imbalance around differentiation and division events as functions of *K* and *t* ≥ 0:

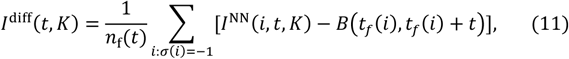

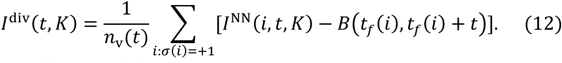

Here, *n*_f_(*t*) and *n*_v_(*t*) are the total number of differentiation and division events that have *I* (*i*,*t* + *t_f_*(*i*)) ≠ NaN, respectively. The indices *i* of differentiating (dividing) cells is denoted as *i*: *σ*(*i*) = −1 (+1) for the sum. Again, the sum in the right hand side skips entries with NaN values.

In Fig. 2i and Extended Data Fig. 2c, we plotted *I*^diff^(*t*,*K*) and *l*^div^(*t*,*K*) as functions of time with *K* = 6. The errors come from fluctuations in *I*^NN^(*i*, *t*, *K*) and *B*(*t*_1_, *t*_2_). We also plot the expected theory line 1 − *e*^-*t/τ*^ assuming a stochastic process for the compensation, where *τ* = 2.1 days and *τ* = 1.0 day were the best fit of the fate-coordination time scale from the fluctuation analyses (Fig. 2d,e and Extended Data Fig. 2a,b). In Extended Data Fig. 3a, we plotted *I*^ditt^(*t*,*K*) as a function of *K* at various time points. The net imbalance quickly saturates at *K* = 4 to 6, indicating that the fate-coordination is occurring within the nearest neighbors.

### Backward tracking of neighbor fate imbalance

For dividing cells with positive G1-reporter signal (cell label *i* with *s_f_*(*i*) = 1), we obtained the timepoints of the cell cycle exit, *t_d_*(*i*). For *t* within *t_b_*(*i*) < *t_d_*(*i*) + *t* ≤ *t_d_*(*i*), we computed the past accumulated imbalance:

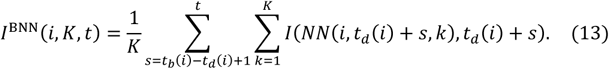

The background of this quantity is the net growth per cell calculated between two time points:

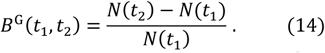

Note that in contrast to *B*(*t*_1_,*t*_2_), the background *B^G^*(*t*_1_,*t*_2_) is zero in the ideal case where there are no fluctuations in division and differentiation rates, irrespective of cell lifetimes. By subtracting this background, we obtained the past accumulated net imbalance:

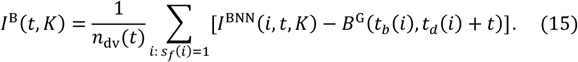

Here, *n_dv_*(*t*) is the number of dividing cells that had positive G1-reporter signal and *I*(*i*,*t* + *t_d_*(*i*)) ≠ NaN. The sum in the right hand side skips entries with NaN values.

In Fig. 3b, we plotted *I^B^*(*t*,*K*) as a function of *t* with *K* = 6. The errors come from the fluctuations of *I*^BNN^(*i*, *t*, *K*) and *B^G^*(*t*_1_,*t*_2_). As a guide to the eye, we co-plotted −*e^t/TC^*, with *r_G_* =1.7 days. This seemingly exponential behavior of *I^B^*(*t*,*K*) indicates that the time it takes from a cell differentiation to one of its neighbor cell exiting G1 phase is stochastic.

### Area growth analysis

We denote the cell area (voronoi area) of cell *i* at time *t* as *A*(*i*, *t*). If cell *i* did not exist at time *t*, then *A*(*i*, *t*) = NaN. The time course of the average cell area for differentiating/dividing cells are (*t* ≤ 0)

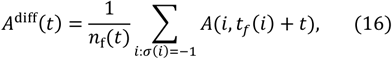

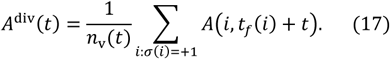

The sum in the right hand side skips entries with NaN values. Similarly, the average of the neighbor cell area of the differentiating/dividing cells are (*t* ≤ 0):

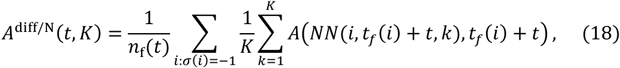

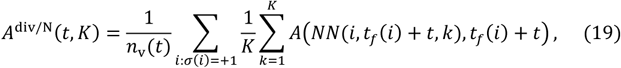

In Fig. 3c, we plotted *A*^diff^(*t*), *A*^diff^(*t*), A^diff*/N*^(*t*,*K* = 6) and *A*^div/N^(*t*,*K* = 6).

**Extended Data Figure 1.**
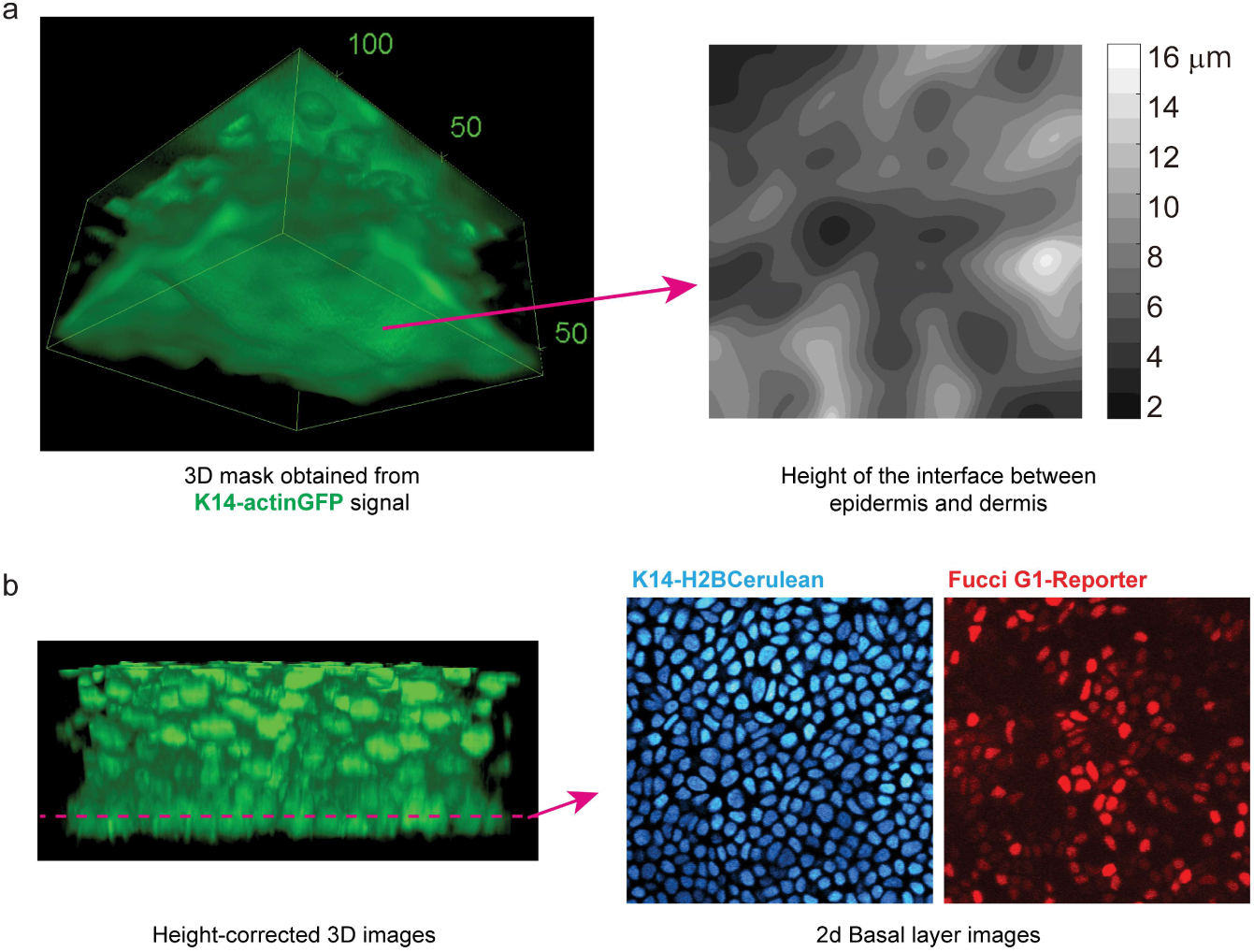
Outline of the image analysis. **a,** Left: example of a 3D structure of the epidermis reconstructed from the K14-actinGFP signal. Raw z-stack image was Gaussian blurred to represent an intact structure. Right: height of the interface between the epidermis (K14-actinGFP positive) and dermis (K14-actinGFP negative) for the left example 3D structure. **b,** Left: original K14-actinGFP image after height correction using information of a. Right: by selecting a z-plane close to the bottom in the height-corrected data, we obtain a 2D image of the basal layer in all channels.

**Extended Data Figure 2.**
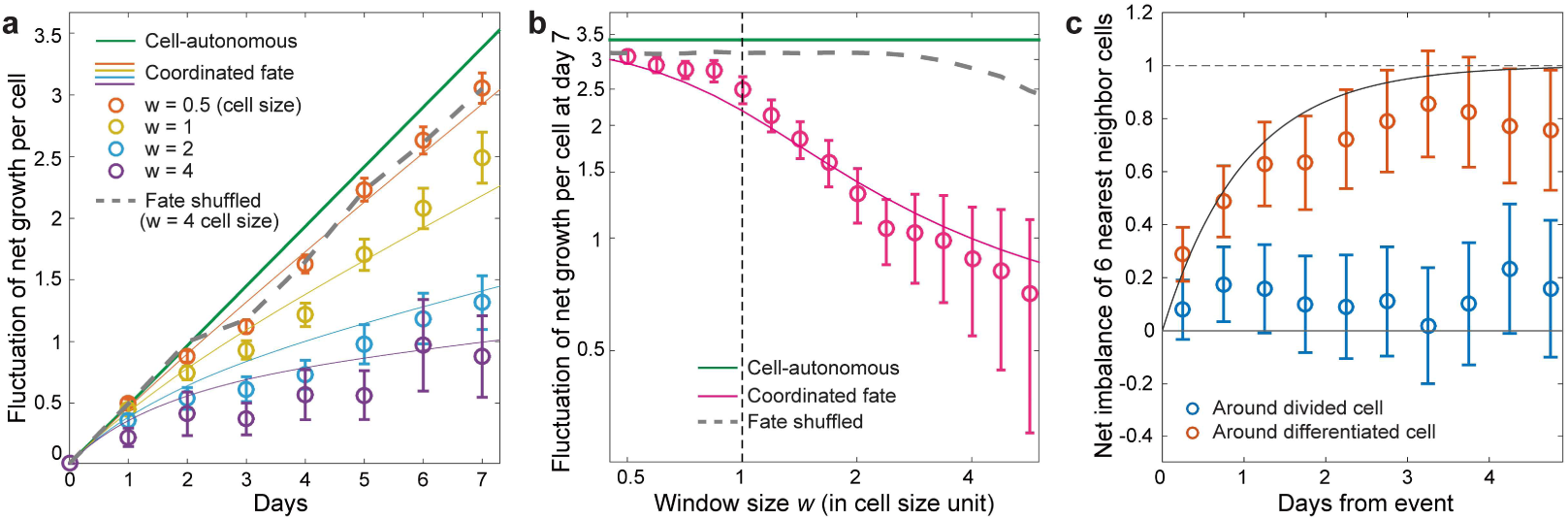
Replicate of fluctuation and neighbor imbalance analyses in a different mouse. **a,b,** Net growth fluctuation analysis, **c**, neighbor net imbalance analysis conducted for a second mouse [321 differentiations and 345 divisions). The qualitative features were the same as the first mouse but with a faster rate (*Λ* = 1.3×10^4^ events day^-1^ mm^-2^). Fitting parameters: *τ* = 1 day, *l* = 4.9 um.

**Extended Data Figure 3.**
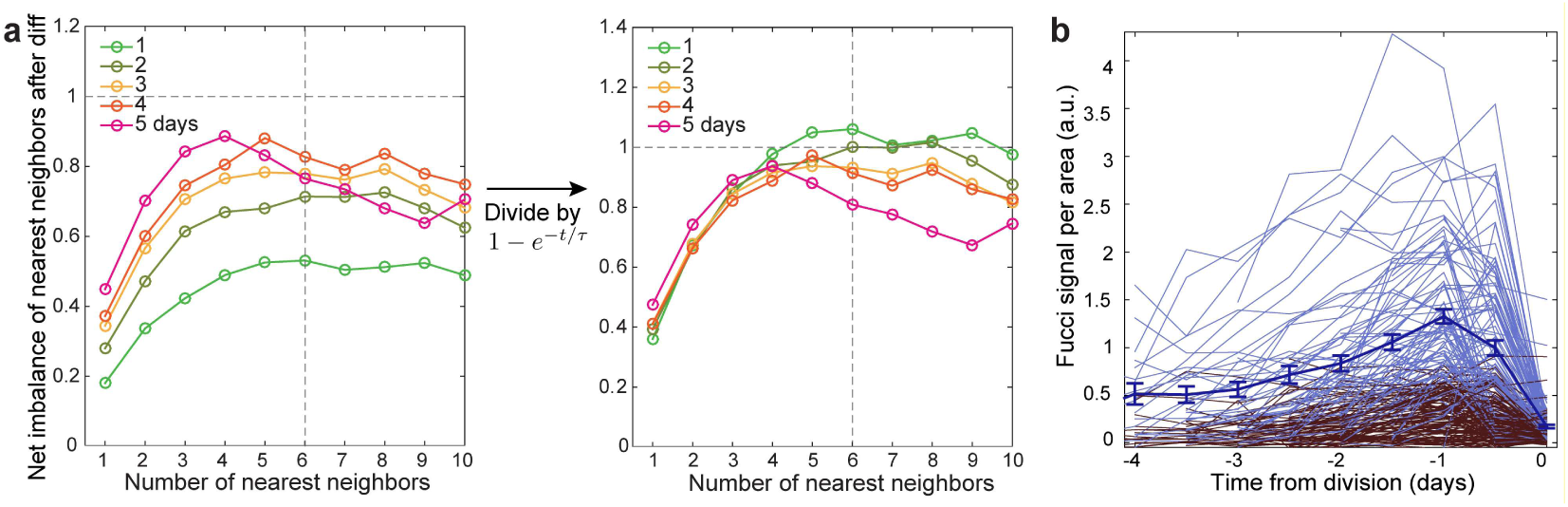
Neighbor number dependence of net imbalance and G1 reporter data. **a,** Left: Net imbalance of nearest neighbor (*I*^diff^(*t*, *K*) defined in Methods) as a function of the number of nearest neighbors. The number of nearest neighbors *K* that is required to saturate *I*^dlff^(*t*, *K*) is the number of relevant neighbors in terms of cell-compensation. We observe saturation at *K* =4 to 6, meaning that the fate-coordination is happening at the nearest neighbor level. Right: We find collapse of data for different *t* when scaled as *I*^diff^(*t*, *K*)/[ 1 − *e ^t/τ^*]. **b,** G1 phase reporter signal per area as a function of the time for each dividing cell. Endpoint of time (*t* = 0) is taken as the cell division time. Blue: G1-reporter positive cells. Dark red: G1-reporter negative cells. Dark blue solid line corresponds to the average of the signal/area for the G1-reporter positive population. Discrimination between G1 signal positive (*s_f_*(*i*) = 1) and negative (*s_f_* (*i*) = 0) was based on a threshold in the maximum value of signal/area. See Methods for detail.

**Extended Data Figure 4.**
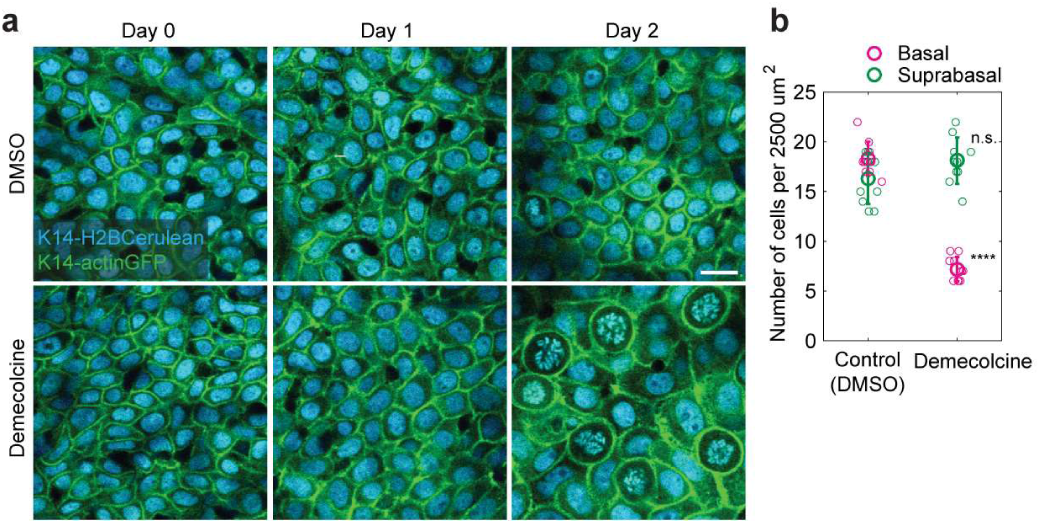
Democolcine cell division blocking data. **a,** Inhibition of stem cell divisions through Democolcine treatment leads to progressive reduction in basal cell density. **b,** Quantification of epidermal basal density. Error bars represent s.d. (10 regions from two mice).

